# Quantitative mitochondrial DNA copy number determination using droplet digital PCR with single cell resolution: a focus on aging and cancer

**DOI:** 10.1101/579789

**Authors:** Ryan O’Hara, Enzo Tedone, Andrew Ludlow, Ejun Huang, Beatrice Arosio, Daniela Mari, Jerry W. Shay

**Affiliations:** Department of Cell Biology, UT Southwestern Medical Center, 5323 Harry Hines Blvd., Dallas TX 75390 U.S.A.; Geriatric Unit, Department of Medical Sciences and Community Health, University of Milan, Via Pace 9, 20122 Milan, Italy; Fondazione Ca′ Granda, IRCCS Ospedale Maggiore Policlinico, Via Francesco Sforza 35, 20122 Milan, Italy.

## Abstract

Mitochondria are involved in a number of diverse cellular functions, including energy production, metabolic regulation, apoptosis, calcium homeostasis, cell proliferation and motility as well as free radical generation. Mitochondrial DNA (mtDNA) is present at hundreds to thousands of copies per cell in a tissue-specific manner. Importantly, mtDNA copy number also varies during aging and disease progression and therefore might be considered as a biomarker that mirrors alterations within the human body. Here we present a new quantitative, highly sensitive droplet digital PCR (ddPCR) method (ddMDM; droplet digital mitochondrial DNA measurement) to measure mtDNA copy number not only from cell populations but also from single cells. Our developed assay can generate data in as little as 3 hours, is optimized for 96-well plates and also allows the direct use of cell lysates without the need for DNA purification or nuclear reference genes. Importantly, we show that ddMDM is able to detect differences between samples whose mtDNA copy number was close enough as to be indistinguishable by other commonly used mtDNA quantitation methods. By utilizing ddMDM, we show quantitative changes in mtDNA content per cell across a wide variety of physiological contexts including cancer progression, cell cycle progression, human T cell activation, and human aging.

## INTRODUCTION

Human mitochondrial DNA (mtDNA) is a circular 16.6 kb genome residing within the mitochondrial matrix that encodes for thirteen protein components of the electron transport chain essential for oxidative phosphorylation (OXPHOS). mtDNA is present at hundreds to thousands of copies per cell, varying widely between cell types and tissues (Robin and Wong 1988; Pierce et al. 1990; Shay et al. 1990). Cellular coordination of mtDNA content is a dynamic and tightly-regulated process (Li et al. 2005; Scarpulla 2006; Wu et al. 2006); however, the mechanisms by which mtDNA copy number is monitored and controlled are not well understood (Moraes 2001; Clay Montier et al. 2009; Klingbeil and Shapiro 2009). Interestingly, alterations in mtDNA levels often accompany key pathophysiological changes during the transition from healthy to diseased states (Butow and Avadhani 2004) and a number of age-related diseases correlate with mtDNA abundance, including cardiovascular disease (Yue et al. 2018), type 2 diabetes (Malik et al. 2009; Monickaraj et al. 2012), cancer (Afrifa et al. 2018), and dementia (Rice et al. 2014; Pyle et al. 2016). Furthermore, mtDNA levels in peripheral blood mononuclear cells (PBMCs) gradually decrease during aging and associate with health status among the elderly (Mengel-From et al. 2014; Wachsmuth et al. 2016) suggesting mtDNA may be a biomarker of biological (not chronological) age and disease exposure (Pieters et al. 2015; Qiu et al. 2015; Tyrka et al. 2015).

The growing relevance of mtDNA as a biomarker highlights the need for a more high-throughput method of mtDNA quantification. The current standard employed in the measurement of mtDNA copy number is quantitative PCR (qPCR). Measurements by qPCR require the use of a reference gene and are often displayed as a ratio of mitochondrial to nuclear DNA; however, qPCR is particularly susceptible to differing PCR efficiencies between target and housekeeping genes, leading to skewing of this ratio (Regier and Frey 2010; Kiselinova et al. 2014). Additionally, the use of a reference gene subjects qPCR results to compounding error, further reducing the sensitivity of qPCR measurement. Recent work has strengthened the utility and flexibility of droplet digital PCR (ddPCR) technology (Ludlow et al. 2014; Huang et al. 2017). ddPCR uses oil emulsion technology to partition samples into thousands of droplets, each representing an independent PCR system. Since all template-containing droplets reach plateau during the PCR step, complications arising from PCR inhibitors and differing PCR efficiencies are thus minimized. The total number of droplets and droplets which fluoresce are counted in a flow cytometry-like fashion to produce a ratio that is then subjected to Poisson distribution, resulting in an absolute quantification of starting template molecules (Hindson et al. 2011; Pinheiro et al. 2012; Robin et al. 2016). Methods to quantify mtDNA copy number by ddPCR using purified genomic DNA have recently been developed (Hindson et al. 2011; Pinheiro et al. 2012; Wachsmuth et al. 2016). However, the time-consuming nature of DNA purification represents a major rate-limiting step in the quantification of nucleic acids (Van Peer et al. 2012) and thus a major hurdle in developing much needed higher-throughput methods for mtDNA copy number evaluation.

Here we present a new quantitative, highly sensitive ddPCR method (ddMDM; droplet digital mitochondrial DNA) to measure mtDNA content per cell equivalent directly from cell lysates or even single cells without the need for DNA purification or nuclear reference genes. The method is optimized for higher-throughput and provides mtDNA measurements in as little as 3 hours from sample collection, making it useful for epidemiological studies. Importantly, ddMDM is more sensitive than previously described assays, reproducibly detecting small differences (<10%) in mtDNA abundance. Finally, by utilizing ddMDM, we show changes in mtDNA content per cell across a wide variety of physiological contexts including cancer progression, cell cycle, human T cell activation, and human aging both *in vitro and in vivo*.

## RESULTS

### ddMDM workflow and experimental design

We optimized the ddMDM workflow (Figure 1A) for accuracy in quantitation, reproducibility, speed and higher-throughput applications (Figure 1A). We initially created 3 sequences (amplicons) representing the mtDNA D-Loop, MT-ND1, and MT-TL1 regions, we made serial dilutions of the amplicons and evaluated the linearity and slopes of three different primers targeting these sequence (Figure 1B, Supplemental Figure 1A-B). We confirmed that measurements taken by using ddPCR accurately reflected absolute molecules per reaction in a nearly 1-to-1 ratio (Figure 1B, Supplemental Figure 1A-B). Because large mtDNA deletions encompassing the D-Loop region have not been previously reported (Bai and Wong 2005), the primer pair targeting the D-Loop region was employed in all subsequent experiments.

**Figure 1.**
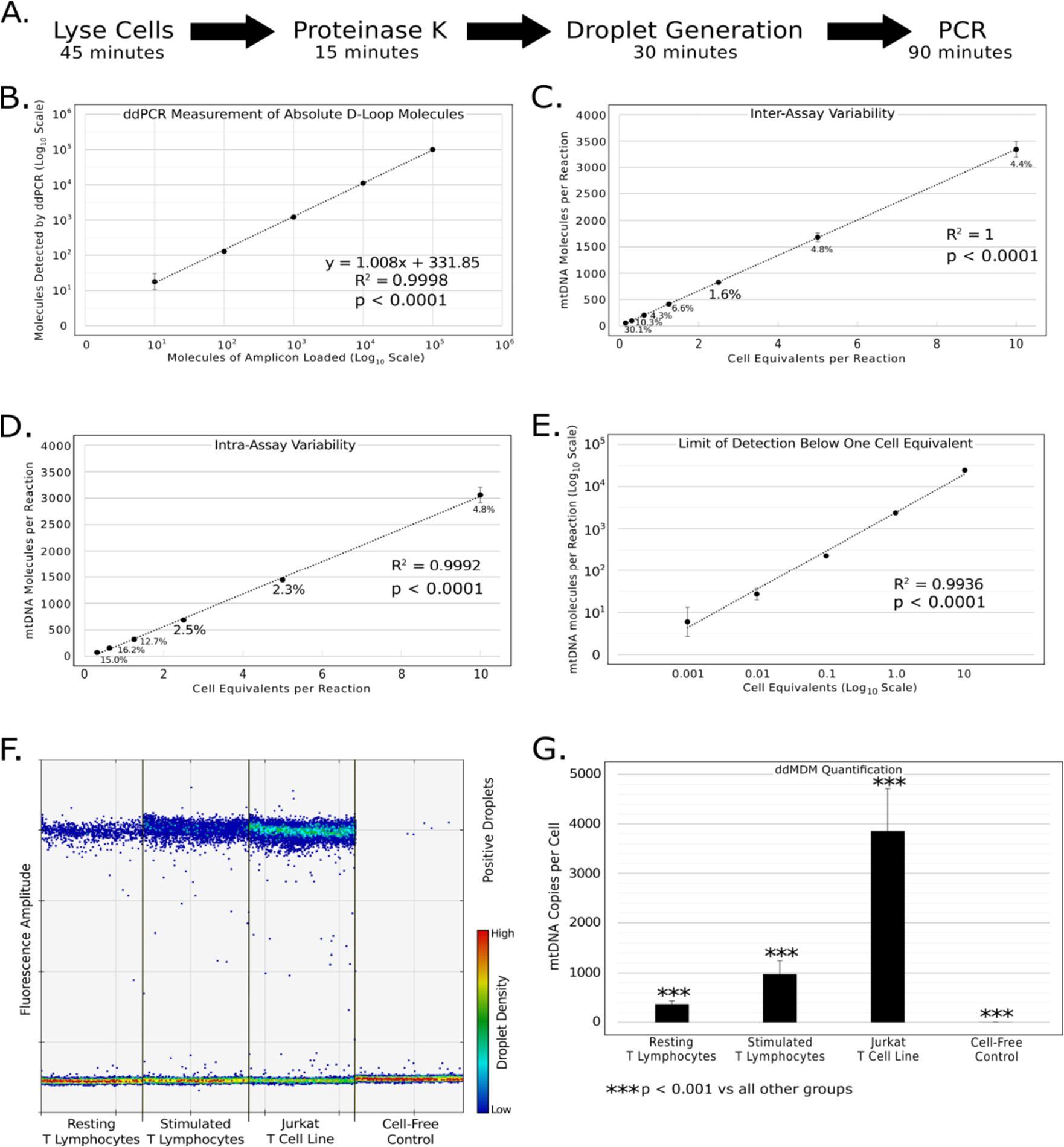
Optimized workflow and validation of method. **A.** General workflow of ddMDM for the quantification of mtDNA on a per cell equivalent basis using ddPCR. **B.** Absolute quantification of D-Loop amplicons by ddPCR in a 10-fold dilution series. Axis are displayed in log_10_ scale. Error bars show the standard deviation of four technical replicates. **C.** Inter-assay variability of ddMDM. Data points represent the mean of means of four biological replicates run in duplicate (eight separate dilution series of early passage BJ fibroblasts’ cell lysates). Adjacent numbers show the coefficient of variation (CV) of the means at that cell equivalents loaded. **D.** Intra-assay variability of ddMDM. Data points represent the average of five technical replicates from a single 2-fold dilution series of early passage BJ fibroblasts. **E.** Limit of detection. 10-fold dilution series in late passage BJ fibroblasts. Data points show the average of two experiments run in triplicate. Error bars display the standard deviation of the six replicates. **F.** ddPCR output showing resting and stimulated T lymphocytes, Jurkat T leukemia cells, and a cell-free (lysis buffer) control. Each lane shows a single ddPCR well, containing an input of 5 cell equivalents. Dots represent emulsion droplets. **G.** Quantification of ddPCR results. mtDNA copy number was calculated per cell equivalent. Error bars show the standard deviation of biological replicates. Significance was determined by ANOVA (Tukey’s multiple comparisons).

Since the results by using serial dilutions of amplicons were highly reproducible (Figure 1B), we next evaluated the reproducibility of the assay when cell lysates rather than artificial amplicons are used. To this aim, we evaluated the intra- and the inter-assay variability by measuring the coefficients of variation (CV) across 2-fold dilution series of BJ fibroblasts cell lysates. We documented very robust reproducibility across 7 serial dilutions with the best reproducibility seen within the range of 800 to 1000 D-Loop molecules per reaction (inter-assay CV=1.6%; intra-assay CV=2.5%; Figure 1C-D). Consistently with other previously published ddPCR methods (Ludlow et al. 2014), this corresponded to ~ 5% of fluorescent (positive) droplets out of the total number of droplets.

We next measured the limit of detection of ddMDM by quantifying mtDNA in a 10-fold dilution series of cell lysates (stimulated T cells), ranging from 10 to 0.001 cell equivalents. Remarkably, ddMDM quantified mtDNA from as little as one hundredth of a cell equivalent (R^2^ = 0.99, Figure 1E). Finally, we tested in a dilution series of cell lysates whether a hydrolysis probe approach (Taq-man) could further improve the accuracy of ddMDM when compared to using EvaGreen but we observed no difference in mtDNA copy number quantitation between the two fluorophores (Supplemental Figure 1C). For this reason the less expensive and simpler to use EvaGreen was employed in all experiments. Together, these results demonstrate that ddMDM reliably quantitates mtDNA copy number by using cell lysates directly. In addition, we quantitated differences in mtDNA content between resting, stimulated, and transformed T cells (Figure 1F). Raw ddPCR output was then transformed into absolute numbers of mtDNA molecules per cell equivalent with a high degree of precision (p < 0.001, Figure 1G).

### ddMDM is more sensitive compared to other commonly employed techniques

Current techniques for assaying mtDNA copy number are often limited in their ability to detect subtle changes in mtDNA levels. As such, it is unknown at what threshold minor perturbations in cellular mtDNA content begin to impact health and disease. We therefore compared ddMDM’s limit of sensitivity to that of other methods by calculating the smallest significant difference that was detectible by each technique. To test this for other techniques, we mixed mitochondrial and genomic amplicons in fixed ratios, simulating samples that differed in mtDNA copy number by set percentages (10% increments in mtDNA) compared to a baseline control. Student’s t-test was used to compare each sample against the baseline control and one-way ANOVA was employed to compare each sample against every other sample. In addition, the standard error of the estimate (σ_est_) was calculated as a measure of the overall variability in each assay. We observed that qPCR could significantly (p < 0.05, Student’s t-test) distinguish between samples whose mtDNA/nDNA copy number ratio differed by at least 50% (Figure 2A), whereas the more stringent ANOVA showed that differences could only be detected at 60% (p < 0.05). The σ_est_ for qPCR detection was 21.3%, reflecting how even moderate deviations in threshold cycles (Ct) can produce large changes in relative abundance calculations.

**Figure 2.**
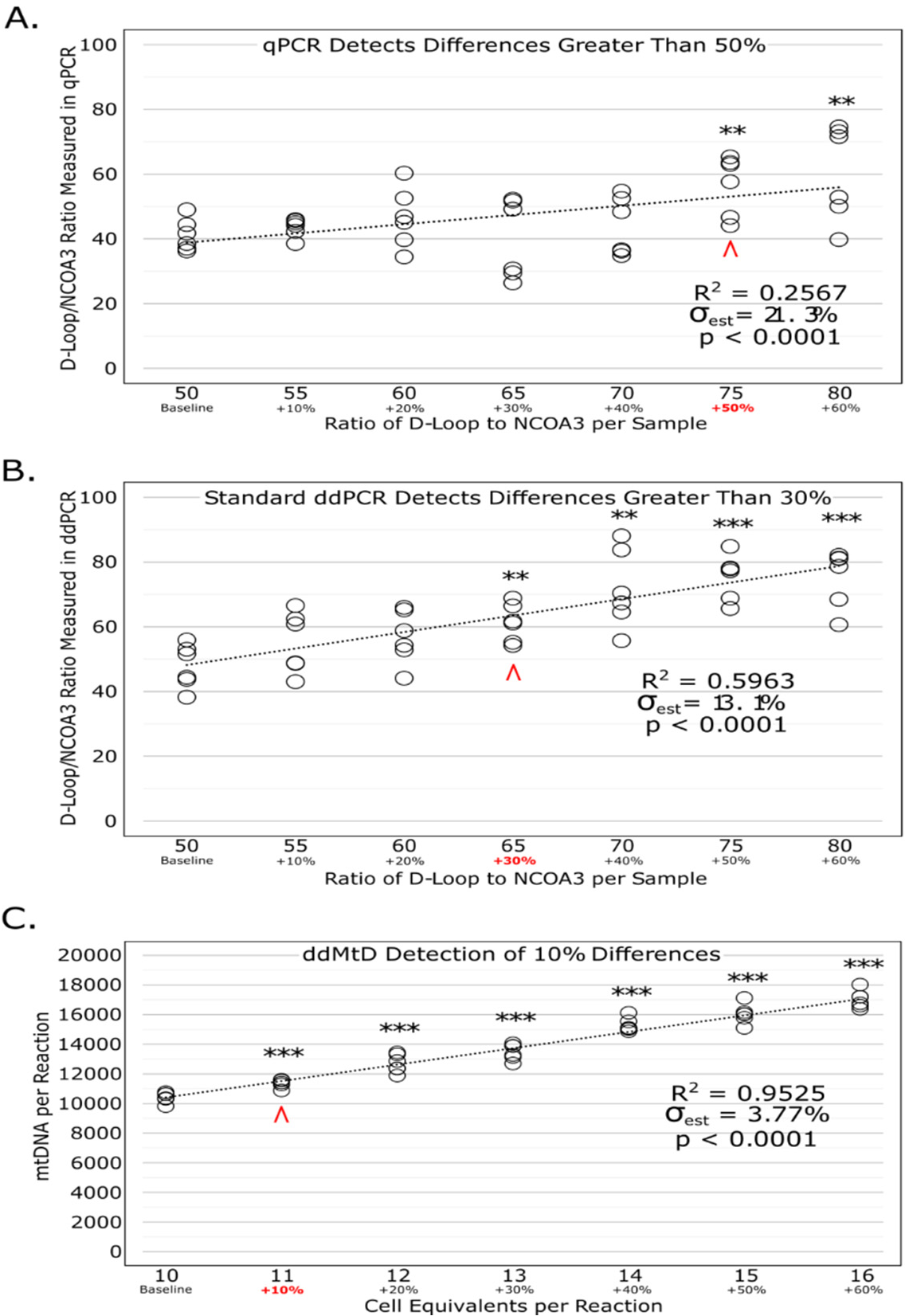
Limit of sensitivity for ddMDM and other previously reported techniques for mtDNA detection. Data points are the result of two separate experiments each performed in triplicate. Asterisks indicate significance by Student’s *t*-test between each sample compared to the baseline control. σ_est_ displays the standard error of the estimate as a percentage, normalized to the population mean. **A.** qPCR determination of D-Loop/NCOA3 ratio using purified amplicons. **B.** ddPCR determination of D-Loop/NCOA3 ratio of the exact same samples employed in Figure 2A. **C.** ddMDM determination of mtDNA molecules (D-Loop) per cell equivalent (stimulated T lymphocytes).

In contrast, ddPCR can detect significant differences (p < 0.05, Student’s t-test) as low as 30% above the baseline, with a σ_est_ of 13.1% (Figure 2B). ANOVA showed that all comparisons with above 33% difference in sample loaded were significant. Importantly, the exact same DNA samples were used for both qPCR and ddPCR experiments, emphasizing that ddPCR use of endpoint rather than threshold fluorescence produce less variable results and quantifications closely resembling the sample ratio that was prepared (Pinheiro et al. 2012; Kiselinova et al. 2014; Zhao et al. 2016).

Finally, we tested the limit of sensitivity for ddMDM using a dilution series of cell lysates from stimulated human T lymphocytes. We found that ddMDM was able to distinguish, with a high degree of significance (p < 0.001, Student’s t-test), all samples from the baseline sample (Figure 2C). Furthermore, ANOVA showed that samples characterized by at least a 9.1% difference in mtDNA content were significantly different. Unlike standard qPCR and ddPCR assays, ddMDM forgoes the use of a nuclear reference gene and thus eliminates the compounded error caused by variability of an additional sample in calculating copy number. The advantage of this is further highlighted by the σ_est_ of only 3.77% (Figure 2C). In summary, we show that ddMDM can detect significant differences between samples whose mtDNA copy number was close enough as to be indistinguishable by other established mtDNA quantitation techniques.

### Detection of mtDNA in a variety of human cells by ddMDM

In order to show the utility and flexibility of ddMDM, we evaluated mtDNA content in various cell types and cellular contexts. Firstly, we examined differential abundance in mtDNA in normal and transformed human cells, having hundreds to thousands of mtDNA copies per cell (Figure 3A). Consistent with previously reported studies, we interpret our results to suggest that mtDNA content varies widely depending on both cell size and cell line specific metabolic profiles (Robin and Wong 1988; Veltri et al. 1990).

**Figure 3.**
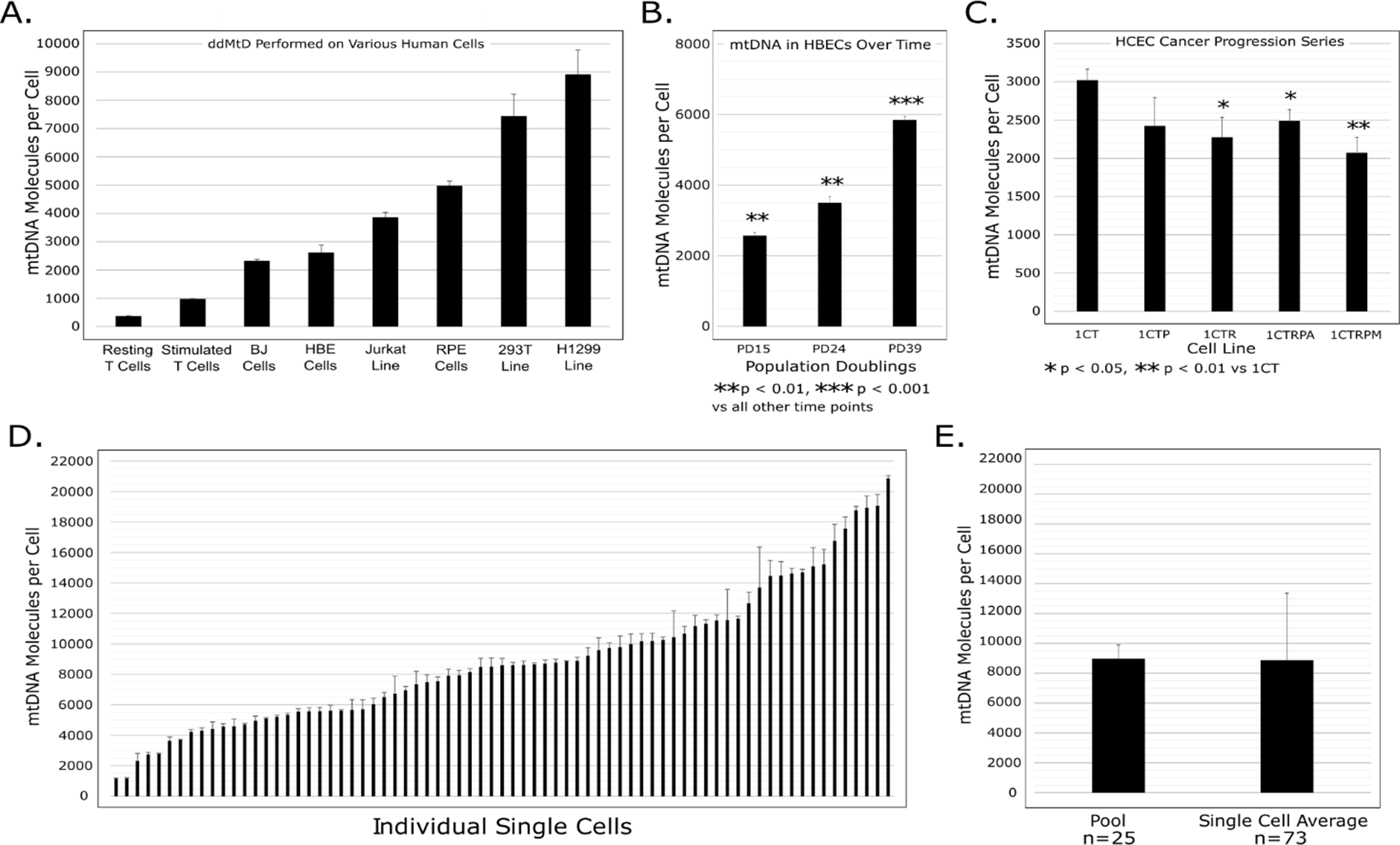
ddMDM quantification of mtDNA in different cell types. **A.** Absolute mtDNA molecules per cell equivalent were measured in different primary and transformed cell lines. **B.** mtDNA levels in primary HBECs at the indicated population doublings. **C.** mtDNA levels in human colonic epithelial cell (HCEC) cancer progression series. Error bars show the standard deviation of biological triplicates. **D.** mtDNA from 73 single H1299 cells were quantified. Error bars show the standard deviation of three technical replicates for each individual cell. **E.** Comparison between averages of H1299 pooled controls and of the 73 single cells.

Changes in mtDNA copy number during aging have been shown to be tissue and cell type specific (Wachsmuth et al. 2016): while mtDNA levels decrease in human peripheral immune cells during aging (Mengel-From et al. 2014) they can increase in other cell types. For example, increases in mtDNA content occur during *in vitro* replicative aging of human bronchial epithelial cells (HBECs) (Figure 3B). Interestingly, the most significant increase in mtDNA copies per cell occurred at around 3-4 population doubling (PD) prior to replicative senescence (reached at PD 42) (Figure 3B) and was accompanied by loss of differentiation potential and a decreased replicative capacity (data not shown).

Metabolic reprograming is one of the key hallmark of cancer (Ward and Thompson 2012) and mtDNA copy number has been reported to correlate with cancer invasiveness and overall patient outcomes (Chen et al. 2016; Hu et al. 2016). In addition, it has been suggested that lower levels of mtDNA in certain cancers may reflect a more stem-like metabolic program and be important in cancer stem cell biology (Lee et al. 2015; Menendez 2015). We used the normal HCEC (human colonic epithelial cells) system with experimentally introduced mutations that commonly occur during colorectal cancer initiation and early progression to examine changes in mtDNA copy number (Eskiocak et al. 2011; Zhang et al. 2015). Specifically, we employed ddMDM to assess how mtDNA abundance varied in response to loss of tumor suppressors (*TP53* and *APC*), oncogenic mutations (*K-rasv12)*, and *TERT, CDK4* and *Myc* overexpression in normal HCECs. We found a significant decrease in mtDNA per cell in the context of K-ras (1CTR and 1CTRPA) and K-ras + Myc (1CTRPM) overexpression (p < 0.05 and p < 0.01 respectively, Figure 3C) suggesting that these cells may be reverting to a more stem-like, undifferentiated state characterized by limited OXPHOS and a preference towards glycolysis (Menendez 2015; Kawada et al. 2017).

### ddMDM for quantitation of mtDNA copy number in single cells

During disease progression, as well as normal aging, key cellular subsets within a mixed cell population often play varying and crucial roles. However, when these peculiar cell subsets are rare, they become diluted and dispersed within the heterogeneous population beyond detection (Mendenhall et al. 2016; Ibrahim-Hashim et al. 2017). In cancer, for example, metabolic reprogramming within tumor subpopulations represents one possible molecular mechanism by which chemotherapy-induced resistance can be acquired (Morandi and Indraccolo 2017). Therefore, in order to further characterize diverse and metabolically distinct cell subpopulations, efficient methods for measuring mtDNA copy number at the single cell level are needed.

We next investigated whether ddMDM was sensitive enough to reliably measure mtDNA copy number in single cells. Briefly, starting with a clonal population, single H1299 human lung cancer cells were plated in 1 μL aliquots onto glass slides and visually confirmed for the presence of exactly one cell by microscopy. Each individual cell was then lysed in a separate tube and 0.3 cell equivalents were employed for the ddMDM assay. Due to the exceptional sensitivity of ddMDM, less than a single-cell equivalent was sufficient to provide an adequate ddPCR signal. This allowed us to perform all single-cell ddMDM in triplicate (from the same individual cell), providing technical replicates for each single cell reaction. We found that mtDNA content varied widely from cell to cell (Figure 3D). Importantly, the average mtDNA content from single cells was nearly identical to that of the pooled cells (mixed population) (Figure 3E), suggesting that our results reflect extreme intercellular differences in mtDNA copy number rather than some type of technical artifact. Additionally, these large differences between single cells cannot be fully explained by cell cycle-dependent variations in mtDNA copy number (Supplemental Figure 1D). Although we observed a significant increase in mtDNA copy number during the transition from G2 to M phase, and a significant decrease during the early G1 phase, these changes were modest when compared to single cell mtDNA heterogeneity (Figure 3D, Supplemental Figure 1D). Thus, this intercellular heterogeneity may be explained, at least in part, by the presence of varying cell sub-populations as previously suggested (Januszyk et al. 2015). Future exploration of mtDNA and metabolic control at the single cell level might reveal a complex and dynamic landscape predicated upon both cell autonomous and microenvironment features. In summary, these results show that ddMDM can detect differences in mtDNA content in single cancer cells from a supposedly homogeneous population.

### mtDNA levels during T cell stimulation decrease with age but healthy centenarians escape this decline

Previously, we identified genes and pathways potentially involved in the process of healthy human aging and longevity through the study of stimulated peripheral blood mononuclear cells (PBMC) from a well characterized population of 114 individuals aged 23-113 years old (Tedone et al. 2019). PBMCs are a heterogeneous cell population mainly consisting of T cells, a major component of human immune responses. T cells remain in a resting or quiescent state when unstimulated, showing no proliferation activity, decreased cell size, and lower metabolic activity. In contrast, upon antigen-specific activation T cells rapidly divide and exhibit dramatic changes in gene expression (Zhao et al. 2014; Tedone et al. 2019) and metabolic features (Dimeloe et al. 2017). Activated T cells initiate immune responses such as discriminating between healthy and abnormal (e.g. infected or cancerous) cells in the body and thus represent a valuable model to study the physiological function of the adaptive immune system. Previous studies have shown a progressive decline of mtDNA copy number in resting PBMC during aging (Kim et al. 2013; Mengel-From et al. 2014; Wachsmuth et al. 2016). However, comparing stimulated PBMC mtDNA levels in different age groups, including centenarians (100+ years old), is an understudied area of research and might provide new insights about the mechanisms driving the age-related impairment of the immune system function. Thus, we stimulated PBMCs from 22 healthy young adults (aged 33±7 years), 22 old (aged 73±6 years), and 17 centenarians (aged 104±4 years) and employed ddMDM to track changes in mtDNA levels over a 15 day (Young) or 10 day (Old and Centenarian) period following stimulation. Since centenarians’ health status has been reported to strongly correlate with their immune cell replicative potential and gene expression (Tedone et al. 2019) we subdivided our centenarians’ study group into two subgroups: healthier centenarians and frail centenarians (Tedone et al. 2019) (Supplemental Table 1).

Consistent with previous studies (D’Souza et al. 2007; Dimeloe et al. 2017), upon stimulation mtDNA levels were transiently up-regulated, generally peaking 3-6 days after stimulation and then slowly declining (Figure 4A). A small but significant dip in mtDNA per cell can be seen at day 1 (422±44 vs 298±85 at days 0 and 1 respectively, p < 0.0001), possibly indicative of activation-induced cell death occurring in a subset of cells (Kabelitz et al. 1993). Unstimulated PBMCs (day 0) showed lower levels of mtDNA, in line with these cells’ quiescent state, and no significant differences were detected between the study groups (Figure 4B). Although previous studies have reported decreased mtDNA in unstimulated PBMCs during aging (Kim et al. 2013; Mengel-From et al. 2014; Wachsmuth et al. 2016), our study found no significant differences.

**Figure 4.**
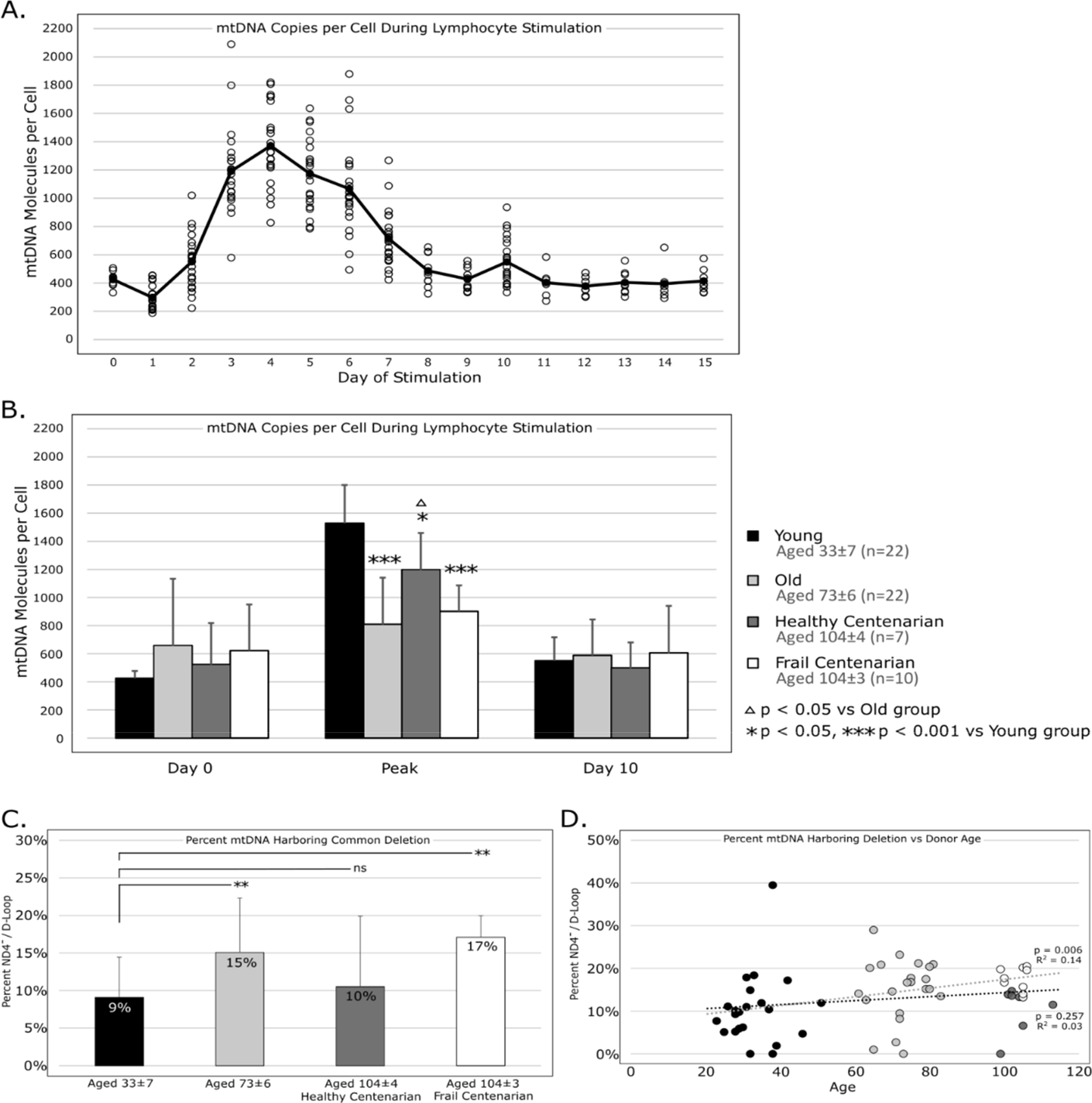
**A.** T lymphocytes from 22 healthy donors were stimulated at day 0 by anti-CD3/CD28 beads and mtDNA per cell equivalent was measured over a 15-day period of stimulation. Open circles represent individual mtDNA per cell from each of the 22 individuals, while the line traces the average of all individuals at each day. **B.** T lymphocytes from 22 healthy young (aged 33 ± 7), 22 healthy old (aged 73 ± 6), 7 healthy centenarian (aged 104 ± 4), and 10 frail centenarian (aged 104 ± 3) individuals were stimulated as described above. mtDNA was measured over the course of the stimulation and the highest point that mtDNA reached for each individual was designated as the “peak” mtDNA of that individual’s stimulation. Shown are averages and standard deviation of the peak mtDNA of each age group. Significance was determined by one-way ANOVA. **C.** ND4 and D-Loop molecule were measured from T cells to show the prevalence of mutated mtDNA (carrying the 4977bp common deletion) to total mtDNA. Error bars show the standard deviation of each group. Significance was determined by ANOVA (Tukey’s multiple comparisons). **D**. The percentage of mtDNA carrying ND4 deletion was plotted versus age for all 61 study participants. Linear regressions were performed for young, old, and frail centenarians (grey line) and young, old, and healthy centenarians (black line).

During stimulation, however, a noticeable impairment in mtDNA biogenesis can be seen with increased age, as measured by the mtDNA copy number peak of each individual over the course of the stimulation period (809±302 vs 1528±272 in old vs young respectively, p < 10^-8^, Figure 4B). This suggests an age-related impairment in PBMCs’ ability to upregulate mtDNA biogenesis in response to antigen-specific activation. Interestingly, we found that mtDNA copy number peak was significantly higher in healthy centenarians as compared to old (aged 73±6) (Figure 4B), suggesting mtDNA biogenesis during stimulation as a potential biomarker of healthy aging.

### ddMDM to quantify common mtDNA deletions: a pilot study in centenarians

Over the course of an individual’s lifespan, mtDNA mutations and deletions accumulate within cells and tissues, in some instances mutant mtDNA propagates at an even faster rate than its wildtype counterparts (Trifunovic et al. 2004; Lakshmanan et al. 2018). It has therefore been suggested that such age-related increases in mtDNA heteroplasmy may contribute to the onset and progression of various diseases (Cortopassi et al. 1992; Chen et al. 2011). The most frequent mtDNA mutation is the “mtDNA common deletion,” a 4977 bp deletion occurring between two 13 bp direct repeats resulting in the disruption of mitochondrial complexes I, IV, and V (Cortopassi and Arnheim 1990; Krishnan et al. 2007; Meissner et al. 2008). In the present study, we adapted ddMDM to multiplex ND4, a gene lying within the deletion, and D-Loop, a region present in all replication-competent mtDNA molecules, such that the proportion of mtDNA molecules carrying the 4977bp common deletion can be calculated as 1 - ND4/D-Loop.

As expected, stimulated PBMCs from young adults displayed a significantly lower prevalence of the “mtDNA common deletion” as compared to both old (aged 73±6) and frail centenarians (Figure 4C). Further, when we performed a linear regression correlating the percentage of mtDNA molecules carrying the common deletion with age in young, old, and frail centenarians we observed a significant age-related decrease in the percentage of intact (unmutated) mtDNA (p = 0.006, Figure 4D). Interestingly, we found no significant difference between young adults and healthy centenarians (Figure 4C) and when we substituted healthy centenarians for frail centenarians and performed the same linear regression, we no longer observed a significant correlation (Figure 4D). Together, these data can be interpreted to show that mtDNA deletions not only accumulate during aging but also that retaining more intact mtDNA appears to be a feature of healthy centenarians and healthy aging. Future investigations are required to explore whether the ability to sustain a higher pool of unmutated mtDNA is a consequence of good health and a relatively disease-free status or whether it promotes optimal health status and it is perhaps causal in some centenarians’ ability to maintain health into extreme age.

## DISCUSSION

The growing number of epidemiological studies exploring the role of mtDNA in health and disease showcases the need for more sensitive and more efficient methods for mtDNA copy number quantification. qPCR is still a major method in the field of mtDNA biology and as a result mtDNA copy number measurements are often confounded by a lack of absolute quantification. The relative quantifications employed by most mtDNA investigations make comparisons between different studies difficult to interpret, often producing conflicting results. Additionally, both PCR efficiency (Regier and Frey 2010; Hu et al. 2016) and the manner in which DNA is isolated (Andreu et al. 2009; Malik et al. 2011) present hurdles to understand the roles of mtDNA copy number during aging and disease progression.

By developing ddMDM, we were able to directly address these key limitations, while providing the added benefit of reduced workflow time, increased sensitivity to detect subtle alterations and quantifications in single cells. With ddMDM we documented changes in mtDNA content during cancer progression, cell cycle progression, cell heterogeneity, T cell activation, and aging. In particular, we observed an age-related reduction in mtDNA copy number in stimulated PBMC from a well characterized human population aged 23-113. Interestingly, healthy centenarians appeared to retain more mtDNA molecules and less major mtDNA deletions, suggesting mtDNA copy number and maintenance during immune cell stimulation is a potential biomarker of healthy aging. In addition, previous studies have shown mtDNA to be a critical player in the NLRP3 inflammation pathway (Zhou et al. 2011; Zhong et al. 2018) and we recently reported involvement of the NLRP3 inflammasome in centenarian longevity and healthy aging (Tedone et al. 2019). Additional studies are required to better understand the contribution that mtDNA plays in chronic inflammation and aging, not only for its role in metabolism but also as a signaling molecule. In fact, cellular mitochondrial content is believed to modulate global transcription (Guantes et al. 2015) but how fluctuations in mtDNA content during aging and disease affect the transcriptome is an area of interest which remains mostly unexplored.

In summary, ddMDM is a novel and rapid (3 hours) method that provides highly quantitative and sensitive mtDNA measurements.

## METHODS

### Cell Culture

Human PBMC were obtained from 61 donors in accordance with the Institutional Review Board (IRB)-approved protocol (certification number STU 042014-016) (Tedone et al. 2019). Subjects affected by cancer, infections or autoimmune diseases or on immunosuppressive treatment at the time of enrollment were excluded from the study. The participant’s age at time of enrollment was defined by birth certificates or identity documents. A trained multidisciplinary staff collected from the enrolled volunteers information regarding their health status together with past and current disease history. Venous blood samples were drawn from the participants under fasting conditions at the same time in the morning. The study protocol was approved by the Ethical Committee of Saint Orsola - Malpighi University Hospital (Bologna, Italy) and written informed consent was obtained from all subjects in accordance with the International Declaration of Helsinki.

PBMCs were isolated by a density gradient centrifugation procedure as previously described (Tedone et al. 2015) and were then cryopreserved at −140 °C pending analysis. Cells were thawed 24 hours prior to stimulation and cultured in RPMI+GlutaMAX-I with 10% fetal bovine serum, 1% penicillin, streptomycin and amphotericin B (Gemini Bioproducts, West Sacramento, CA). After 24h the cell suspensions were transferred into a new flask and PBMCs were stimulated by adding Dynabeads Human T-Activator CD3/CD28 (Life Technologies) in a 1:1 cell-to-bead ratio. After 72 hours of stimulation Dynabeads were removed using a magnet and cells were cultured for up to 15 days after stimulation. The percentage of live cells was determined every day by trypan blue exclusion using a TC20 Automated Cell Counter (BioRad). Cell density was maintained at 1.0×10^6^ cells/mL.

Jurkat (T leukemia) cells were cultured in RPMI+GlutaMAX-I with 10% fetal bovine serum, 1% penicillin, streptomycin and amphotericin B (Gemini Bioproducts, West Sacramento, CA) and maintained at 1.0×10^6^ cells/mL. H1299 (non-small cell lung adenocarcinoma), 293T (human embryonic kidney), RPE (retinal pigment epithelial), and various populations of normal human BJ (foreskin fibroblasts) cells were cultured in ambient oxygen, 5% CO_2_ and maintained in a 4:1 ratio of DMEM (Dulbecco’s modified Eagle’s medium, GE Healthcare, Logan, UT) to Medium 199 containing 10% calf serum (HyClone, Logan, UT, USA). Primary HBECs (human bronchial epithelial cells) were grown in 2% oxygen using a 1:1 mixture of bronchial epithelial basal medium (BEBM; Lonza) and DMEM and supplemented with SingleQuot (Lonza) and 1% PSA. Primary dermal foreskin fibroblasts (early passage BJ fibroblasts) were isolated and cultured as previously described (Ramirez et al. 2003).

The HCEC (human colonic epithelial cell) cancer progression series lines (1CT, 1CTP, 1CTR, 1CTRPA, 1CTRPM) were grown as described above. Initially, diploid HCECs lines were established by TERT (T) and CDK4 (C) immortalization. Then, additional lines bearing combinations of commonly found mutations during colorectal cancer initiation and early development were established. (A: APC shRNA stable knockdown and truncated APC A1309 overexpression, M: cMyc overexpression, P: TP53 shRNA stable knockdown, R: mutant Kras^v12^ overexpression) (Eskiocak et al. 2011; Zhang et al. 2015).

### Cell lysis procedure for mtDNA quantification

Pellets containing 1×10^5^ cells were collected and stored at −80°C pending analysis. Cell pellets were lysed on ice using 40 µl of NP-40 lysis buffer (Figure 1A) and vortexed every 15 minutes for a total of 45 minutes. Once lysed, 10 µl of lysate (containing 2.5×10^4^ cell equivalents) was added to 9 µl of TNES buffer (10mM Tris pH 8.0, 100mM NaCl, 10mM EDTA, 1% SDS) and 1 µl of proteinase K (Invitrogen 20mg/mL). Proteinase K digestion was performed at 50°C for 15 minutes, followed by 100°C for 10 minutes, then cooled to 12°C. Following proteinase K treatment, samples were diluted with ddH_2_O 1:250 to obtain the final concentration of 5 cell equivalents per µl. After dilution, samples were prepared for ddPCR. ddPCR is sensitive to oversaturation of template molecules and optimal sample dilution will vary between cell types, dependent on mtDNA content per cell.

### ddPCR by using cell lysates (ddMDM)

Each 20 µl ddPCR reaction contained a final concentration of 1X EvaGreen ddPCR Supermix (Bio-Rad), 100 nM D-Loop forward, 100 nM D-Loop reverse primer, and 1 µl of sample (1-5 cell equivalents). Samples were partitioned into droplets using a QX100 droplet generator (Bio-Rad) according to the manufacturer’s instructions and the emulsions (approximately 40 µl) were transferred to a 96-well PCR plate (twin-tec, Eppendorf) and sealed with foil (Thermo Scientific, AB0757). Following droplet formation, PCR was performed at 95°C for 5 minutes, 40 cycles of 95°C, 54°C, and 72°C for 30 seconds each, then held at 12°C. The ramp rate between all steps was 2.5°C/second. After PCR, fluorescence was measured using a QX100 droplet reader (Bio-Rad), detecting, on average, 17-20 thousand droplets per sample. The threshold for positive droplets was determined by the software’s analysis of droplet clustering across all samples and confirmed manually by comparison to a negative, cell-free control (Figure 1B). The final output given by the software was a concentration of starting template molecules per µl and could be converted to molecules per 20 µl reaction (the original pre-emulsion volume containing 1-5 cell equivalents). Finally, mtDNA per reaction was normalized to obtain copies of mtDNA per cell equivalent (ddMDM) (Figure 1C).

For quantifications of mtDNA expressed as molecules of mtDNA per single-copy nuclear gene (D-Loop/NCOA3 ratio), two reactions per sample were prepared: one containing 100nM D-Loop primers, the other containing 100nM NCOA3 primers. After PCR the number of mtDNA copies per diploid genome was calculated as previously described (Bai and Wong 2005).

### Total genomic DNA extraction

Extraction and purification of total genomic DNA was performed as previously described (D’Souza et al. 2007). Briefly, 2×10^5^ cells were pelleted and resuspended in 250 µl of SDS lysis buffer. Samples were briefly vortexed then boiled for 10 minutes. Samples were cooled to room temperature and treated with 2.5 µl of RNAse A (10mg/mL, Qiagen) at 37°C for 2 hours. Next, 2.5 µl of proteinase K (10mg/mL) was added and samples were incubated at 55°C overnight. The next day samples were boiled for 10 minutes, followed by DNA precipitation by adding 0.1 volumes of 3M sodium acetate and 2 volumes of 100% ethanol at −20°C overnight. Centrifugation was performed at 4°C at 1500g for 15 minutes. Pellets were washed once in 70% ethanol and resuspended in 50 µl ddH_2_0. DNA concentration of samples was measured by Qbit High Sensitivity DNA kit (Thermo Fisher Scientific).

### ddPCR by using purified genomic DNA

mtDNA measurements from genomic DNA by using ddPCR were performed as described (Wachsmuth et al. 2016). Briefly, total genomic DNA was isolated and ddPCR was performed as described above. Absolute mtDNA copies per ddPCR reaction were normalized to the single-copy nuclear gene NCOA3. In order to generate a dilution curve for normalized results, mtDNA was measured in serial dilutions of total genomic DNA and then normalized to the nDNA measured from 0.1ng of the same DNA.

### qPCR

All qPCR reactions were performed as previously described (Bai and Wong 2005) and consisted of 1 µl of sample (1 ng total DNA), 1X Ssofast EvaGreen Supermix (BioRad), 500 nM forward, and 500 nM reverse primer to a final volume of 20 µl. qPCR was performed using a Lightcycler 480 (Roche) at 95°C for 5 minutes, 40 cycles of 95°C for 15 seconds and 60°C for 60 seconds, followed by melting curve analysis. Samples measured in qPCR represent the average of three technical replicates, unless otherwise indicated. In order to convert threshold cycles to absolute molecules, standard curves of known amounts of purified D-Loop and NCOA3 amplicons were run concurrently with qPCR samples. Exponential regressions created from these standard curves were then used to calculate mtDNA and nDNA per ng of DNA employed. Dilution curves of normalized results were calculated as described above.

### Preparation of purified amplicons

One nuclear and three mitochondrial primer pairs were chosen, targeting NCOA3 (Bai and Wong 2005), the D-Loop region (Bai and Wong 2005), MT-ND1 (Kim et al. 2013), and MT-TL1 (Bai and Wong 2005). These primers were confirmed for specificity by both agarose gel and by qPCR melting curve analysis. PCR amplicons of these four primer pairs were then purified from 1% agarose gel using Qiagen Gel Extraction kit and concentrations were measured with Qbit High Sensitivity DNA kit (Thermo Fisher Scientific). Each amplicon was then diluted to 1×10^9^ molecules per µl, aliquoted, and stored at −20°C.

### Single-cell mtDNA quantification

48 hours prior to collection, cells were cultured at low density in order to avoid potential contact-induced inhibition at the single cell level. After trypsinization, PBS was added to bring the cell density to around 1×10^6^ cells/mL. 1×10^5^ cell pellets were collected to be simultaneously run alongside single-cell samples as pooled control samples. The remaining cell suspension was further diluted in PBS +0.1% FBS to roughly 2-4 cells per µl. This suspension was aliquoted onto a glass slide in 1 µl droplets (figure 4B). Each individual aliquot of cell suspension was confirmed by microscope for the presence of exactly one cell before being transferred to PCR tubes containing 0.05 µl proteinase K (25 mg/mL, 1.25 ng final), 0.45 µl TNES buffer, 1 µl NP-40 lysis buffer, and 7.5 µl MS2 RNA in ddH_2_O (32 pg/µl, 240 pg final). MS2 RNA helped to prevent excessive quantities of DNA from sticking to tubes during single cell reactions. The final 10 µl reaction was run in a thermocycler at 50°C for 30 minutes, 100°C for 10 minutes, then cooled to 12°C for lysis and proteinase K treatment. 15µl of ddH_2_O was then added and the sample mixed thoroughly by pipetting. 7.5 µl of diluted sample (corresponding to 0.3 cell equivalents) was then used to perform ddPCR reactions as described above, with the exception of differing sample volumes. All single-cell ddMDM reactions shown were run in triplicate.

### Statistical analysis

Statistical analyses were performed using Student’s *t*-test (unpaired, two-tailed), ANOVA (Tukey’s multiple comparisons), or linear regression where appropriate. Significance was denoted as follows: * *p* ≤ 0.05; ** *p* ≤ 0.01; *** *p* ≤ 0.001; **** *p* ≤ 0.0001.

## ACKNOWLEDGEMENTS

This work was supported by the National Institutes of Health [AG01228], the Harold Simmons National Cancer Institute Designated Comprehensive Cancer Center Support Grant; the Southland Financial Corporation Distinguished Chair in Geriatric Research [JWS]. This work was performed in laboratories constructed with support from National Institutes of Health [grant C06RR30414].

---

**Supplemental Figure 1.**
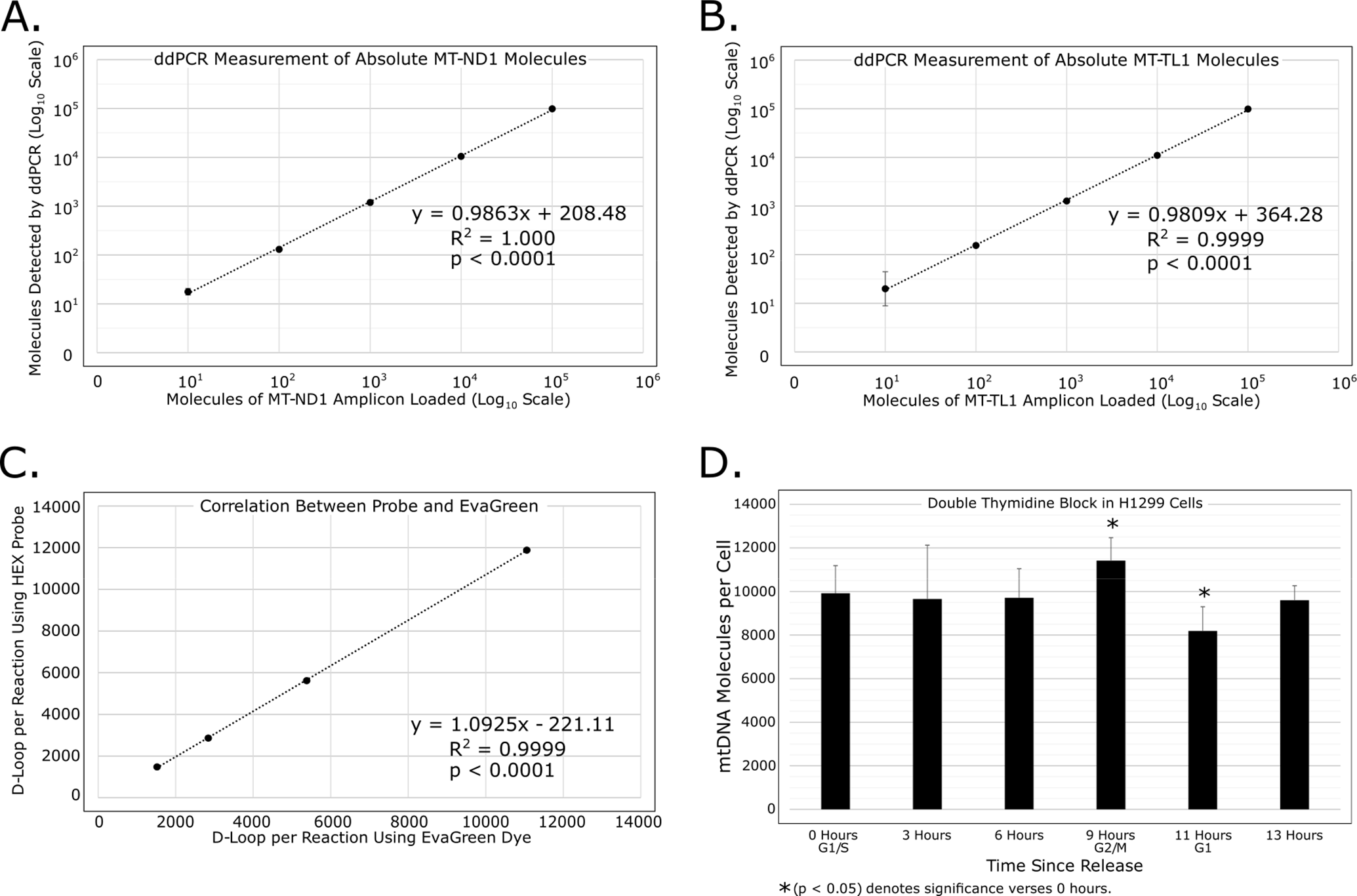
Absolute quantification of additional mtDNA amplicons using ddPCR were measured in 10-fold dilution series. Axis are displayed in log_10_ scale. Error bars show the standard deviation of four technical replicates. **A.** MT-ND1, NADH Dehydrogenase Subunit 1. **B.** MT-TL1, Mitochondrially-encoded tRNA Leucine 1 (UUA/G). **C.** Correlation between use of Taq-man probe and use of EvaGreen dye in the quantification of D-Loop molecules, performed in a 2-fold dilution series from stimulated T lymphocytes. **D.** mtDNA levels in H1299 cells at the indicated time points post-release from G1/S. Error bars represent the standard deviation of six biological replicates (two separate synchronizations run in triplicate).

**Supplemental Table 1.** Health status assessment of centenarians. Disease count was determined by evaluating the presence of the following diseases: Acute myocardial infraction, stroke, angina, hypertension, COPD, dementia, depression, diabetes, thyroid dysfunction, arthrosis, chronic liver diseases and chronic kidney diseases.

| Health status assessment | Healthier Centenarians<br>(n=7) | Frail Centenarians<br>(n=10) |
| --- | --- | --- |
| Age, years, mean ± S.D. | 105.6 ± 4.2 | 104.4 ± 2.0 |
| Gender, females | 5/7 | 7/10 |
| Smokers, % | 0 | 0 |
| Body Mass Index (BMI), mean ± S.D. | 24.1 ± 2.0 | 21.9 ± 2.7 |
| Disease count per individual, mean ± S.D. | 1.7 ± 0.5 * | 4.8 ± 1.6 |

\* $p < 0.05$

